# Endogenous glucocorticoids moderate the gastric inflammatory response to *Helicobacter* infection and protect from autoimmunity

**DOI:** 10.1101/2025.07.03.662976

**Authors:** Sara R. Druffner, Benjamin C. Duncan, Maeve T. Morris, Jordan L. Pascoe, Tyler M. Abner, Salik Hussain, M Blanca Piazuelo, Richard M. Peek, Melody Zhang, Richard J. DiPaolo, Jonathan T. Busada

**Author notes:** **Corresponding author:** Jonathan T. Busada, WVU School of Medicine, Microbiology, Immunology and Cell Biology, 64 Medical Center Drive, P.O. Box 9177, Morgantown, WV 26506. Authors contributed equally (Sara R. Druffner, Benjamin C. Duncan). **Disclosures:** The authors have declared that no conflict of interest exists. **Author Contributions:** SRD, BCD, MTM, JLP, TMA, and JTB performed all experiments and analyzed data. SH performed the digital PCR. MBP performed histopathology scoring. MZ and RJD performed the autoantibody screening. RMP and RJD provided critical expertise, assisting with data interpretation. SRD, BCD, and JTB drafted the manuscript. All authors reviewed and approved the final manuscript.

## Abstract

**Background and Aims:** Immune responses to infection must balance pathogen clearance with minimizing tissue damage and autoimmunity. Chronic gastric inflammation caused by *H. pylori* damages the gastric mucosa and promotes carcinogenesis. Glucocorticoids are immunoregulatory hormones that limit immune activation in the stomach. This study aimed to determine how endogenous glucocorticoids regulate the gastric immune response to *Helicobacter* infection and their impact on preneoplastic lesion development.

**Methods:** We examined the role of endogenous glucocorticoids in shaping the gastric immune response to *Helicobacter felis* colonization. Gastric immune cell infiltration, atrophy, metaplasia, and preneoplastic lesion development were evaluated in adrenal-intact control mice and adrenalectomized (ADX) mice. Auto-reactive IgG antibodies were assessed using a mouse self-antigen array and by measuring their binding to healthy gastric tissue.

**Results:** Loss of endogenous glucocorticoids led to significantly increased *H. felis-*induced gastric T cell infiltration and proinflammatory cytokine expression compared to intact-infected controls. While all intact mice maintained chronic infection for up to 12 months, nearly all ADX mice eradicated *H. felis* within 2–3 weeks. Despite bacterial clearance, ADX mice continued to exhibit chronic gastric inflammation and developed dysplasia. Autoantibody profiling showed that both intact and ADX groups generated self-reactive IgG during active infection. However, only ADX mice sustained autoantibody production following bacterial eradication.

**Conclusions:** Endogenous glucocorticoids attenuate gastric inflammation during *Helicobacter* infection, supporting bacterial persistence while maintaining immune tolerance. These findings suggest that heightened immune responses to *H. pylori* may trigger autoimmune gastritis (AIG) development, which can persist after *H. pylori* clearance and continue to drive gastric cancer risk.

## Introduction

Gastric cancer is the fifth most common cancer and the fifth leading cause of cancer-related deaths worldwide ^1^. Chronic inflammation is a driver of gastric carcinogenesis and is most commonly associated with *Helicobacter pylori* infection. However, recent studies have also linked autoimmune gastritis (AIG) to an increased risk of gastric adenocarcinoma ^2–5^. Although *H. pylori* infection and AIG are distinct disease processes, severe pathogen-induced inflammation is known to trigger autoimmunity through mechanisms such as host tissue damage, co-presentation of microbial and self-antigens, and heightened proinflammatory cytokine production that promotes bystander activation of self-reactive T cells ^6–8^. *H. pylori* infection has long been suspected of contributing to autoimmunity and clinical studies have detected anti-parietal cell antibodies in infected individuals ^9^. Nevertheless, the connection between *H. pylori* and AIG remains poorly defined, and further investigation is needed to determine whether *H. pylori* can initiate or exacerbate autoimmune responses in the stomach.

The risk of gastric adenocarcinoma is closely tied to the nature and intensity of the host immune response. Strong inflammatory responses promote the development of preneoplastic lesions, whereas tolerogenic responses limit tissue damage and are protective ^10^. Glucocorticoids are anti-inflammatory steroid hormones produced by the adrenal glands that signal through the glucocorticoid receptor to maintain immune homeostasis ^11^. They function by suppressing proinflammatory cytokine production, reducing antigen presentation and co-stimulation, and promoting regulatory T cell differentiation ^12^. Endogenous glucocorticoids play a critical role in maintaining gastric immune balance. We have previously reported that bilateral adrenalectomy (ADX), which depletes circulating glucocorticoids, leads to spontaneous gastric inflammation, atrophic gastritis, and metaplasia within the gastric corpus ^13–15^. However, their role in regulating gastric inflammation during *H. pylori* infection remains unclear.

Although *H. pylori* prevalence has declined in the United States over recent decades, the incidence of early-onset gastric cancer—particularly in young women—has increased ^16, 17^. These cancers often arise in the absence of detectable *H. pylori* infection and are frequently associated with AIG. Yet, the underlying etiology of these cases, including whether *H. pylori* infection previously occurred and contributed to cancer initiation, remains unknown.

In this study, we investigated the role of endogenous glucocorticoids in regulating gastric inflammation in response to *Helicobacter* infection. We colonized ADX mice and adrenal-intact controls (hereafter referred to as “intact”) with *Helicobacter felis*. ADX mice exhibited enhanced gastric T cell responses and significantly increased expression of Th1-associated proinflammatory cytokines. While infection persisted in intact mice, ADX mice cleared *H. felis*. Both groups developed autoantibodies in response to infection, but ADX mice exhibited greater antigen diversity and higher titers of IgG antibodies targeting gastric antigens. However, despite bacterial clearance, ADX mice continued to display atrophic gastritis, metaplasia, and progressed to dysplasia. Together, these findings demonstrate that endogenous glucocorticoids moderate the gastric immune response to *Helicobacter* infection. While this immune suppression facilitated bacterial persistence, it also protected against autoimmunity. Moreover, our results suggest that *Helicobacter* infection may initiate autoimmune gastritis that continues to promote neoplastic progression even after bacterial eradication.

## Material and Methods

### Animal Care and Treatment

All mouse studies were approved by the Animal Care and Use Committee at West Virginia University. C57BL/6J female mice were purchased from the Jackson Laboratories. Only female mice were used in this study as we have previously shown that androgens compensate for loss of glucocorticoids in male mice ^14^. Mice were administered standard chow and water *ab libitum* and maintained in a temperature- and humidity-controlled room with standard 12-hour light/dark cycles. For all experiments, adrenalectomy surgeries were performed at 8 weeks of age. Following adrenalectomy, mice were sustained on 0.9% saline drinking water to maintain ionic homeostasis. Within one week of surgery, ADX mice and age-matched adrenal-intact controls were either mock infected or infected with *H. felis* by oral gavage. Mock mice received 500μL sterile brucella broth, and infected mice were inoculated with 500 μL of brucella broth containing 10^9^ CFU two times 24 hours apart.

### Bacterial Preparation

*Helicobacter felis* was grown on tryptic soy agar plates (BD Biosciences) with 5% defibrinated sheep blood (Hemostat Labs, Dixon, CA) and 10 μg/mL vancomycin (Thermo Fisher Scientific, Waltham, MA) under microaerophilic conditions at 37°C for two days. The bacteria were then harvested and transferred to Brucella broth (RPI, Mount Prospect, IL) containing 5% fetal bovine serum (R&D Systems, Minneapolis, MN) and 10 μg/mL vancomycin and grown overnight at 37°C under microaerophilic conditions with agitation. Bacteria were centrifuged and resuspended in fresh Brucella broth without antibiotics before spectrophotometry and mouse infection.

### Antibiotic Treatments

Antibiotics were dissolved in ethanol and then diluted in water to a final concentration of 0.1% ethanol, 15 mg/kg amoxicillin (MP Biomedicals, Solon, OH), 12 mg/kg metronidazole (Thermo Fisher Scientific), and 2 mg/kg omeprazole (Thermo Fisher Scientific). Mice were administered antibiotic cocktail twice daily for seven consecutive days by oral gavage. Elimination of infection was confirmed by PCR on fecal DNA.

### Tissue Preparation

Mice were euthanized by cervical dislocation without anesthesia. Stomachs were removed and opened along the greater curvature and washed in phosphate-buffered saline to remove gastric contents. Biopsies were collected and immediately snap-frozen in liquid nitrogen for RNA and DNA isolation. Tissue strips were collected from the greater and lesser curvatures and fixed overnight in 4% paraformaldehyde at 4°C. Strips were either cryopreserved in 30% sucrose and embedded in optimal cutting temperature media or transferred into 70% ethanol and submitted to the histology core at West Virginia University for routine processing, embedding, sectioning, and H&E staining. The remaining corpus was dissociated into a single cell suspension for flow cytometry as described below. We previously reported that ADX induces spontaneous gastric inflammation and metaplasia ^15^. This inflammation is limited to that gastric corpus lesser curvature. Therefore, we avoided the lesser curvature in our analysis to avoid the spontaneous inflammation triggered by ADX.

### Histology

Standard methods were used when performing immunostaining. 5μm stomach cryosections were incubated with anti H+/K+ ATPase (clone 1H9, MBL Life Sciences), MIST1 (Cell Signaling Technologies), CD45 (clone 104 Biolegend), or CD44v9 (Cosmo Bio) for 1 hour at room temperature or overnight at 4°C. Sections were then incubated in secondary antibodies for 1 hour at room temperature. Where indicated, fluorescence-conjugated *Griffonia simplicifolia* lectin (GSII, ThermoFisher) was added with the secondary antibodies. Sections were mounted with Vectastain mounting media containing 4’,6-diamidino-2-phenylindole (Vector Laboratories). Images were obtained using a Zeiss 710 confocal laser-scanning microscope (Carl-Zeiss) and running Zen Black (Carl-Zeiss) imaging software. Parietal cells and chief cells were quantitated as previously described ^15^ using confocal micrographs captured with a ×20 microscope objective. Cells were counted using the ImageJ count tool (National Institutes of Health). Cells that stained positive with anti-H^+^-K^+^ antibodies were identified as parietal cells and cells that stained positive with anti-MIST1 antibodies and were GSII negative were identified as mature chief cells. Cell counts were normalized to image area to determine the cell number per 100 μm^2^. Images that contained gastric glands cut longitudinally were selected for counting.

### RNA isolation and qRT-PCR

RNA was extracted in TRIzol (ThermoFisher) and precipitated from the aqueous phase using 1.5 volume of 100% ethanol. The mixture was transferred to an RNA isolation column (Omega Bio-Tek) and the remaining steps were followed according to the manufacturer’s recommendations. Reverse transcription followed by qPCR was performed in the same reaction using the Universal Probes One-Step PCR kit (Bio-Rad Laboratories) and the TaqMan primers *listed* in Table 1, normalized to *Ppib.* Fold change is in relation to vehicle controls.

### Flow Cytometry

Harvested tissue was diced and washed in Hanks Balanced Salt Solution without Ca^2+^ or Mg^2+^ containing 5 mM HEPES, 5 mM EDTA, and 5% FBS at 37°C for 20 minutes. Tissue pieces were then washed briefly in Hanks Balanced Salt Solution with Ca^2+^ or Mg^2+^ and then digested in 1mg/ml collagenase (Worthington) for 30 minutes at 37°C. After digestion, tissue fragments were pushed through a 100μM strainer and then rinsed through a 40μM strainer before debris was removed through an Optiprep (Serumwerk) density gradient. Fc receptors were blocked with TruStain (Biolegend), then stained with primary antibodies for 20 minutes on ice. Actinomycin D (A1310, Invitrogen) was used to label dead cells. Samples were analyzed on a Cytek Aurora spectral flow cytometer (Cytek Biosciences). Flow cytometry analysis was performed using Cytobank (Beckman Coulter).

### Autoantibody Testing

Serum was collected from mice and sent to the Genomics and Microarray Core Facility at UT Southwestern Medical Center where samples were screened for IgG and IgM autoantibodies. Heatmap was made in Morepheus and is plotted as fold change from the vehicles. A fold change of two was used as the minimum cutoff for including the antigen in the heatmap.

For autoantibody immunostaining, sera were from mock-infected and *H. felis* infected mice and used to probe tissue sections of healthy gastric corpus. Anti-mouse IgG were used as secondary antibodies. Sections from three biological replications were probed with sera from each mouse. Immunofluorescence images were collected with an ECHO Revolve microscope (BICO, Gothenburg, Sweden). To quantify autoantibody staining intensity, images were analyzed using FIJI (National Institutes of Health). Images were split into single channels and converted to grey scale. IgG mean fluorescence intensity (MFI) was measured for the entire and normalized by tissue area. The fluorescent intensity range was 0=black to 255=white.

### Statistical Analysis

All error bars are ± SD of the mean. The sample size for each experiment is indicated in the figure legends. Experiments were repeated a minimum of two times. Data normality was assessed using the Kolmogorov-Smirnov test. Datasets with Gaussian distribution were assessed by one-way ANOVA with the post hoc Tukey’s test. Nonparametric datasets were assessed by Kruskal-Wallis test followed by a Dunn’s t-test. *H. felis* clearance was analyzed using Kaplan-Meier survival curves and compared between groups using the Mantel-Cox test. In parallel, infection rates over time were analyzed using two-way ANOVA with Sidak’s multiple comparisons test. Statistical analysis was performed by GraphPad Prism 10 software. Statistical significance was set at *P* ≤ 0.05. Specific *P* values are listed in the figure legends.

## Results

### ADX increases the T cell response to H. felis infection

We previously reported that endogenous glucocorticoids are key regulators of gastric inflammation and that ADX induced spontaneous inflammation within the gastric corpus, lesser curvature ^18^. To investigate how endogenous glucocorticoids impact gastric inflammation during *Helicobacter* infection, intact and ADX mice were colonized with *H. felis* and assessed two months post-infection. Immunostaining for the pan-leukocyte marker CD45 revealed sparse tissue-resident submucosal and intraepithelial leukocytes in mock-infected controls (Figure 1A). In contrast, *H. felis* colonization stimulated robust leukocyte infiltration in both intact and ADX mice. Flow cytometric analysis revealed a similar proportion of overall leukocytes within the gastric corpus of infected intact and ADX mice (Figure 1B). Similarly, myeloid cell infiltration remained relatively similar between infected intact and ADX mice (Figure 1C). Interestingly, ADX induced a marked shift in the lymphocyte compartment. While *H. felis* colonization of intact mice led to a significant increase of B and T cells (Figure 1D), B cell infiltration was significantly blunted in ADX mice. In contrast, ADX significantly enhanced gastric T cell infiltration (Figure 1D). Interestingly, *H. felis* colonization triggered expansion of regulatory T cells in intact mice, while the proportion of regulatory T cells remained unchanged in ADX mice. Next, we examined the expression of the proinflammatory cytokines *Ifng, Tnf,* and *Il1b* in the gastric corpus. These cytokines remained unchanged in ADX mock-infected controls but were significantly increased in ADX-infected mice compared to intact infected controls (Figure 1E). These results demonstrate that endogenous glucocorticoids moderate gastric T cell activation during *H. felis* infection and suggest that disruption of glucocorticoid signaling may lead to a hyperactive T cell response.

**Figure 1:**
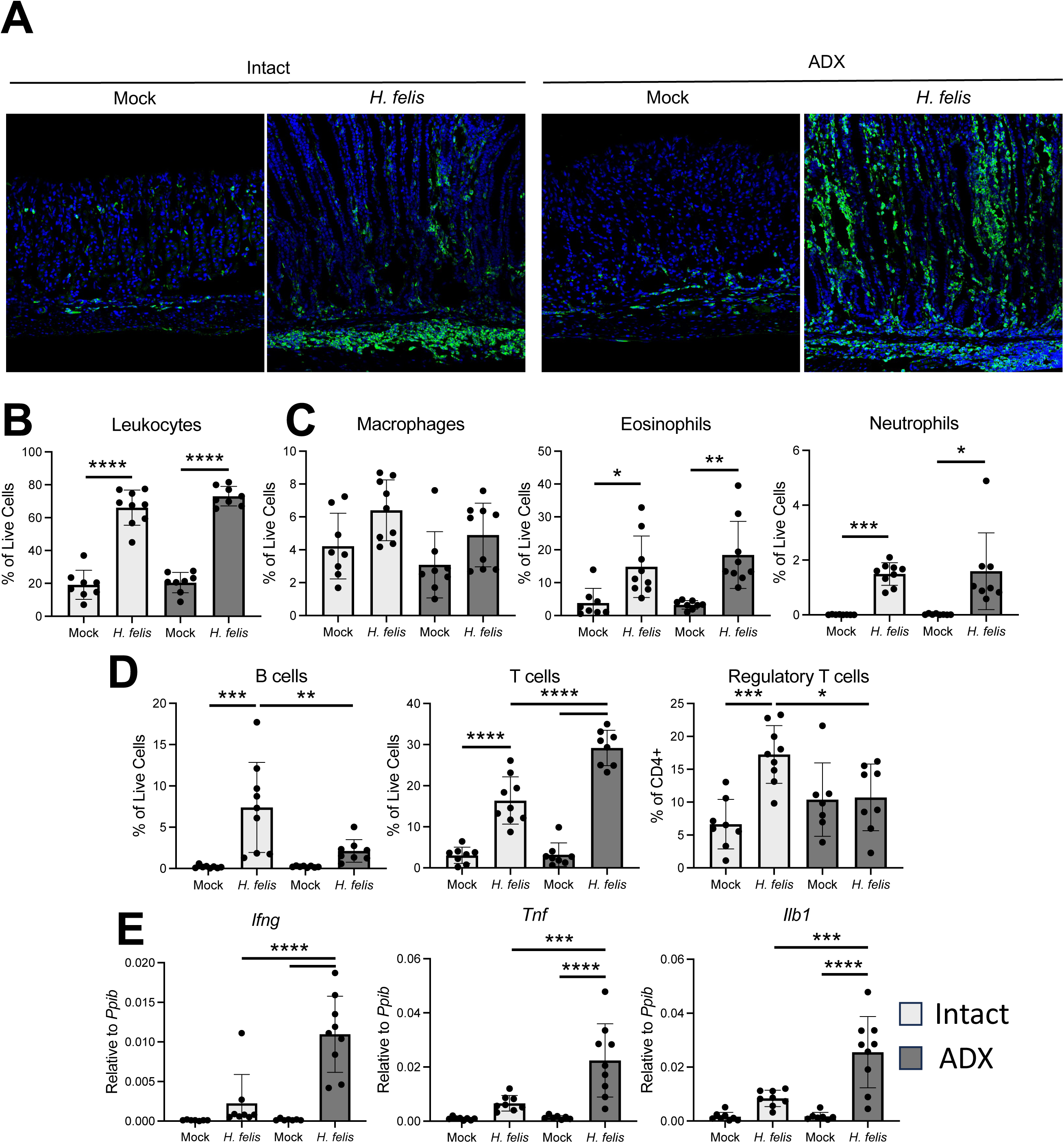
Adrenalectomy increases gastric T cell infiltration during *H. felis* infection. (A) Representative immunofluorescent staining of CD45+ immune cells (green) in the gastric corpus of intact and ADX mice 2 months after mock infection or *H. felis* colonization. Nuclei are counterstained with DAPI (blue). (B-D) Flow cytometry quantification of (B) total infiltrating gastric leukocytes, (C) myeloid compartment (macrophages, eosinophils, and neutrophils), (D) lymphoid compartment (B cells, T cells, and regulatory T cells) in the gastric corpus. (E) qRT-PCR of the indicated proinflammatory cytokines using RNA isolated from the gastric corpus. Data are presented as mean ± SD; \**p* < 0.05, **p < 0.01, \*\*\**p* < 0.001, ****p < 0.0001. n≥7.

### ADX does not significantly alter Helicobacter-driven gastric atrophy but promotes pyloric metaplasia development

To determine how glucocorticoid depletion influences gastric atrophy and metaplasia, key features of *Helicobacter* infection, we examined histological changes in ADX mice. Given that ADX alone induces inflammation, atrophy, and metaplasia primarily in the lesser curvature of the gastric corpus ^15^, we focused our analysis on the greater curvature. Histological analysis of H&E-stained sections and immunostaining for mucous neck, parietal, and chief cell lineages revealed that ADX mock-infected stomachs were largely comparable to intact controls (Figure 2A-B). *H. felis* infection induced significant parietal and chief cell atrophy and mucous neck cell hyperplasia in both intact and ADX mice (Figure 2A-C), although oxyntic atrophy was slightly more severe in the stomachs of ADX mice (Figure 2C). Next, CD44v9 immunostaining was used to label pyloric metaplasia development ^19, 20^. Expression was not detected within the corpus of intact and ADX mock-infected controls (Figure 2D). In contrast, *H. felis* colonization drove widespread CD44v9 expression in both intact and ADX stomachs with positive cells found in the base of nearly every gastric unit. However, CD44v9 expression was qualitatively more intense and widespread in ADX mice. *H. felis* infection significantly upregulated the PM-associated genes *Aqp5, Gkn3*, and *Wfdc2* in both intact and ADX mice, with expression levels significantly higher in the ADX-infected group (Figure 2E). These findings indicate that *H. felis* infection drives complete parietal and chief cell atrophy in both intact and ADX mice. However, ADX mice exhibited enhanced PM development in response to infection, suggesting a role for glucocorticoids in modulating metaplasia progression.

**Figure 2:**
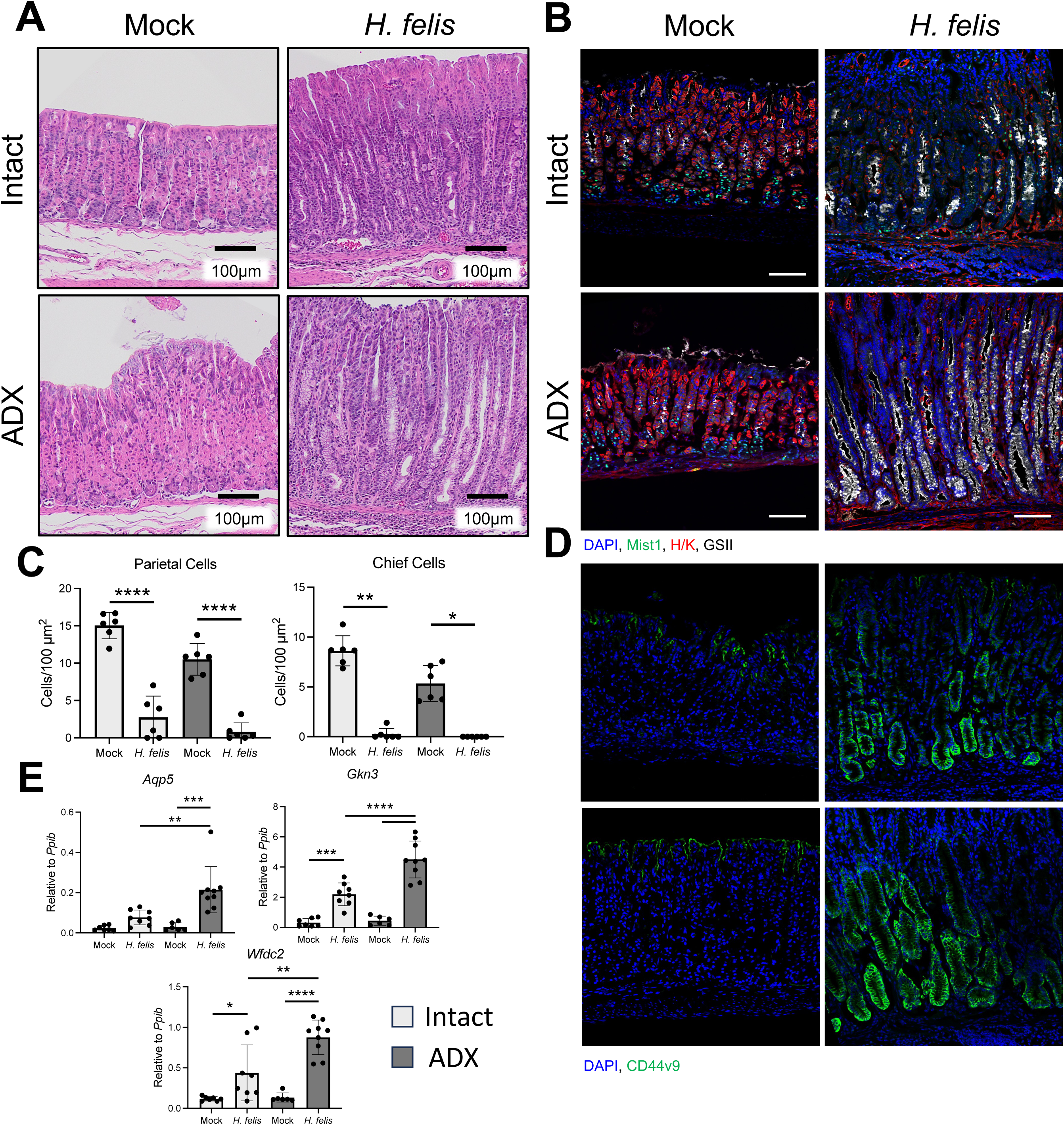
Endogenous glucocorticoids protect from pyloric metaplasia development during *H. felis* infection. Representative micrographs of (A) H&E-stained sections or (B) immunostaining of H+/K+ ATPase (red, parietal cells), MIST1 (green, chief cells), GSII lectin (white, mucous neck cells), and DAPI (nuclei, blue) of the gastric mucosa from intact and ADX mice 2 months post *H. felis* colonization. (C) Quantification of chief and parietal cell loss (D) Immunofluorescent staining of the metaplastic marker CD44v9 (green) in the gastric corpus. (E) qRT-PCR of the metaplasia transcripts *Aqp5, Gkn3,* and *Wfdc2*. Data are presented as mean ± SD; \**p* < 0.05, **p < 0.01, \*\*\**p* < 0.001, ****p < 0.0001. n≥6.

### Glucocorticoid signaling promotes H. felis persistence

To confirm *H. felis* colonization, we performed conventional PCR on DNA extracted from the gastric corpus and antrum. Notably, while *H. felis* DNA was detected in 95% of intact mouse stomachs, it was only present in 33% of ADX mouse stomachs (Figure 3A). We reasoned that conventional PCR may not be sensitive enough to detect low level *H. felis* colonization. Therefore, we performed digital PCR, which is extremely sensitive to low abundance DNA. Digital PCR detected *H. felis* DNA in all stomach samples from intact mice, whereas no *H. felis* DNA was detectable in the stomachs of ADX mice (Figure 3B), suggesting that ADX mice clear *H. felis* infection. To monitor colonization over time, we performed longitudinal PCR analysis of fecal DNA. At one-week post-colonization, *H. felis* DNA was detectable in fecal samples from all mice, confirming successful colonization (Figure 3C). However, by eight weeks post-infection, *H. felis* remained detectable in 9 of 10 intact mice but declined significantly in ADX mice, with only 2 of 10 testing positive (Figure 3C). These findings suggest that endogenous glucocorticoids are critical for sustaining *H. felis* colonization. In the absence of glucocorticoid signaling, *H. felis* is progressively eradicated, indicating that host-derived glucocorticoids promote bacterial persistence.

**Figure 3:**
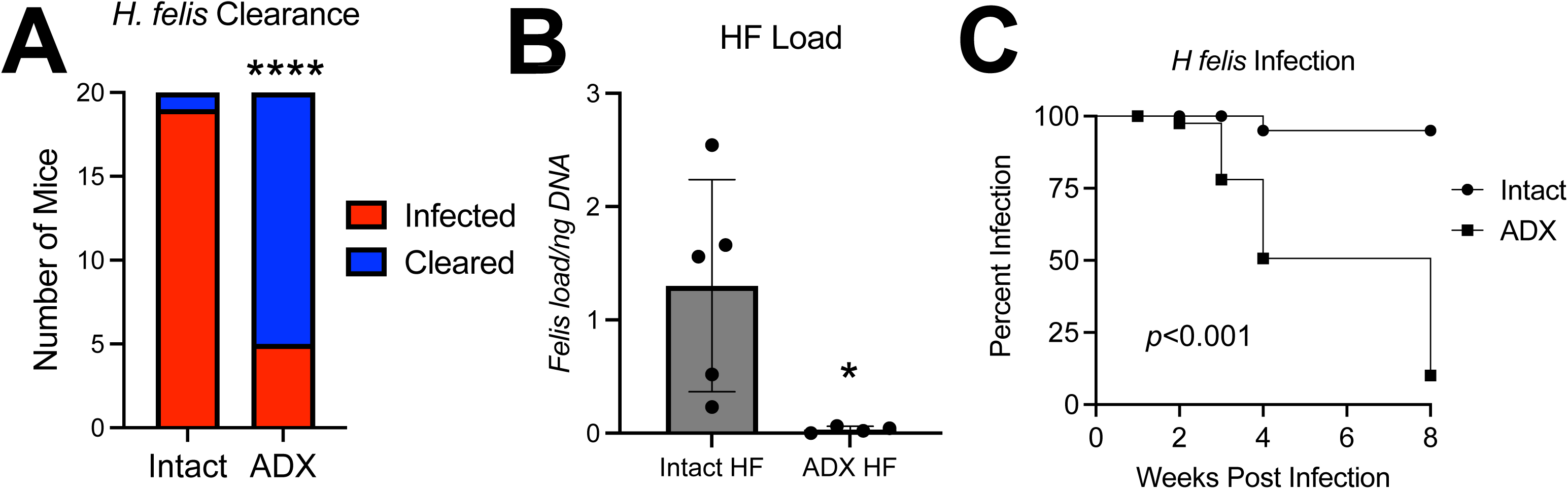
Endogenous glucocorticoid promotes *H. felis* persistence. (A) Graph depicting the number of mice with active *H. felis* infection (red) versus cleared infections (blue) sham vs adrenalectomized mice. (B) Digital PCR of *H. felis* in gastric corpus biopsies n≥4. (C) Kaplan-Meier curve depicting spontaneous *H. felis* eradication. Infection confirmed by PCR of fecal DNA and gastric biopsies. \**p* < 0.05, \*\*\**p* < 0.001, ****p < 0.0001. n≥15.

### Endogenous glucocorticoids are required for inflammation resolution and mucosal recovery following H. felis eradication

While ADX mice clear *H. felis* infection, they continue to exhibit severe gastric inflammation (Figure 1), atrophy, and metaplasia (Figure 2). To further explore this, we colonized intact and ADX mice with *H. felis.* After two months, mice were treated with antibiotics (ABX) to eradicate the infection, and their stomachs were analyzed one-month post-treatment (Figure 4A). One month after ABX treatment, flow cytometric analysis revealed that gastric inflammation had resolved in intact mice, whereas it persisted in ADX mice (Figure 4B). However, T cell infiltration remained significantly elevated in ADX mice despite the absence of *H. felis*. Consistent with the resolution of inflammation, the gastric mucosa of intact mice largely recovered following eradication, while gastric atrophy persisted in ADX mice (Figure 4C-D). Quantification of intact mice demonstrated that parietal cell numbers returned to baseline post-eradication. However, while chief cell numbers increased, they remained significantly lower than in mock-infected controls (Figure 4E). In contrast, *H. felis* eradication did not significantly improve parietal or chief cell counts in ADX mice. Similarly, PM resolved in intact mice following eradication, as shown by the absence of CD44v9 immunostaining and the reduced expression of *Aqp5, Gkn3,* and *Wfdc2* (Figure 4F-G). However, ADX mice continued to exhibit widespread CD44v9 expression and significant elevation of PM marker transcripts. These results demonstrate that ADX mice exhibit persistent gastric inflammation and tissue pathology following *H. felis* eradication, suggesting that disruption of endogenous glucocorticoid signaling may promote the development of self-reactive immune responses.

**Figure 4:**
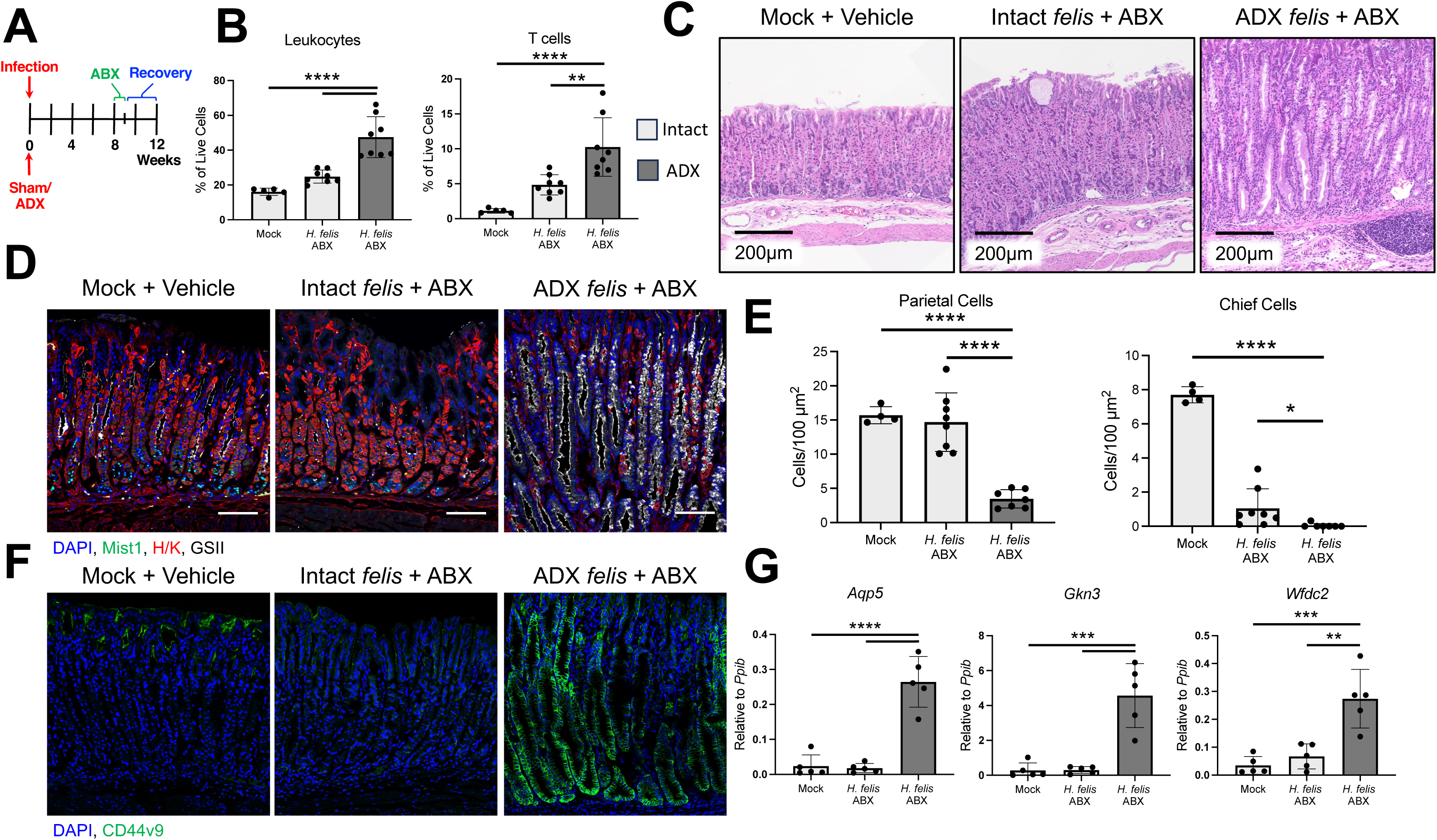
Endogenous glucocorticoids promote gastric epithelial recovery following *H. felis* eradication. (A) Antibiotic treatment schematic. (B) Flow cytometry quantification of gastric leukocytes and T cells within that gastric corpus in antibiotic treated intact and ADX. Representative micrographs of (C) H&E-stained sections or (D) immunostaining of H+/K+ ATPase (red, parietal cells), MIST1 (green, chief cells), GSII lectin (white, mucous neck cells), and DAPI (nuclei, blue) of the gastric mucosa from antibiotic treated sham and adrenalectomized female mice 2 months post *H. felis* colonization. (E) Quantification of chief and parietal cell loss. (F) Immunofluorescent staining of the metaplastic marker CD44v9 (green) in the gastric corpus. (G) qRT-PCR of the metaplasia transcripts *Aqp5, Gkn3,* and *Wfdc2*. Data are presented as mean ± SD; \**p* < 0.05, **p < 0.01, \*\*\**p* < 0.001, ****p < 0.0001. n≥5.

### Despite eradicating infection, preneoplastic progression persists in ADX mice

To investigate the long-term implications of infection on preneoplastic lesion development, we assessed intact and ADX mice 12 months after *H. felis* colonization. PCR analysis confirmed that all intact mice maintained active infection, while no *H. felis* DNA was detected in ADX-infected mice (Figure 5A). Histological analysis revealed no notable abnormalities in the stomachs of intact and ADX mock-infected controls (Figure 5B). Blinded pathological scoring of H&E sections from intact infected mice revealed severe chronic inflammation (Figure 5C), along with significant parietal and chief cell atrophy (Figure 5D). In contrast, ADX mice displayed a heterogenous phenotype. All ADX-infected mice displayed persistent chronic gastric inflammation (Figure 5C). However, parietal and chief cell atrophy remained elevated in 45% and 83% of mice, respectively (Figure 5D). Finally, pathologist evaluation of dysplasia and carcinoma revealed that 60% of intact mice developed dysplasia and 10% developed carcinoma, while 45% of ADX mice developed dysplasia (Figure 5E). These findings demonstrate that, despite clearance of *H. felis*, ADX mice continue to exhibit chronic gastric inflammation with a subset developing preneoplastic lesions.

**Figure 5:**
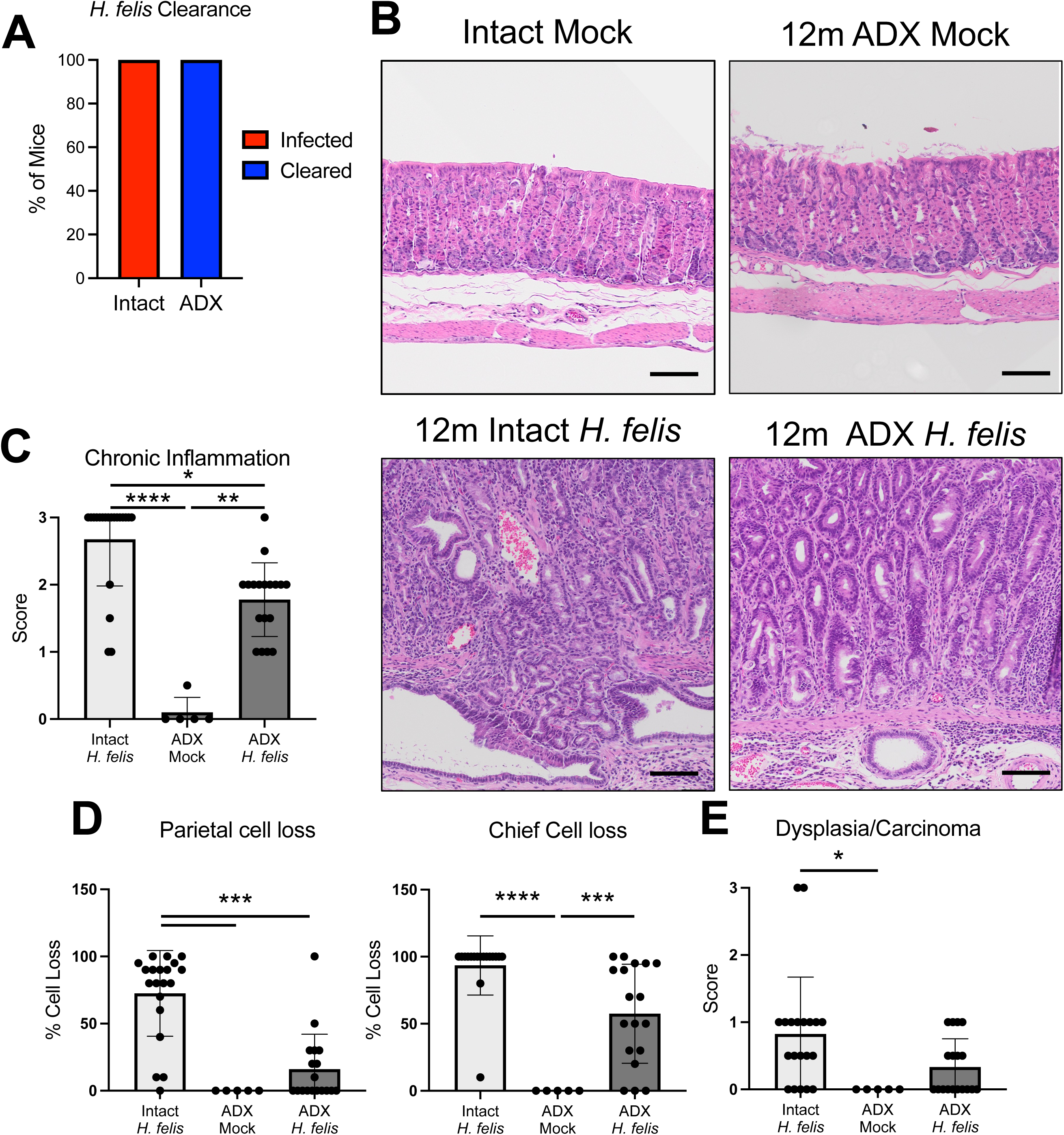
ADX mice develop dysplasia, despite eradicating *H. felis*. (A) Graph depicting the number of mice with active *H. felis* infection (red) versus cleared infections (blue) sham vs adrenalectomized mice. (B) Representative micrographs of H&E-stained sections of mock-infected and *H felis* infected intact and ADX mice 12 months post colonization. (C) Blinded pathologist scoring of the indicated gastric pathologies. Data are presented as mean ± SD; \**p* < 0.05, **p < 0.01, \*\*\**p* < 0.001, ****p < 0.0001. n=5 for ADX-mock and n≥18 for infected groups.

### Endogenous glucocorticoids support peripheral tolerance during H. felis infection

The persistence of chronic gastric inflammation in ADX mice despite *H. felis* clearance suggests that loss of endogenous glucocorticoids may promote autoimmune reaction. Therefore, we collected sera from intact mock-infected, intact *H. felis-*infected, and ADX-infected mice two months post-colonization and screen for self-reactive IgG antibodies. Self-reactive IgG were detected for a wide variety of self-antigens in both intact and ADX-infected groups (Figure 6A). However, IgG antibody binding to gastric intrinsic factor was detected exclusively in ADX mice. To further assess autoreactivity, we collected sera from intact and ADX mice two months after *H. felis* colonization, then the mice were treated with antibiotics to eradicate *H. felis* and sera was again collected one month after eradication (Figure 6B). These sera were used to immunostain gastric corpus sections collected from uninfected mice to evaluate self-reactive IgG binding (Figure 6C). Minimal signal was detected in sections probed with sera from mock-infected controls (Figure 6D). Sera from intact-infected mice showed binding to multiple cell types within the gastric corpus, including clear staining of parietal and mucous neck cells. In contrast, sera from ADX-infected mice exhibited broader reactivity, staining parietal, chief, and mucous neck cells, and producing strong nuclear signals suggestive of anti-DNA IgG (Figure 6D). Quantification of the mean fluorescent intensity confirmed significantly higher levels of self-reactive IgG in the sera from ADX-infected mice (Figure 6F). Finally, sera collected one month after *H. felis* eradication demonstrated a significant reduction of self-reactive IgG in the sera of intact mice (Figure 6E-F). Whereas self-reactive IgG levels remained significantly elevated in the sera of ADX mice. Together, these results demonstrate that *H. felis* infection leads to the activation of self-reactive lymphocytes, and that loss of endogenous glucocorticoid signaling exacerbates the breakdown of peripheral tolerance, increasing both the magnitude and diversity of self-reactive IgG responses.

**Figure 6:**
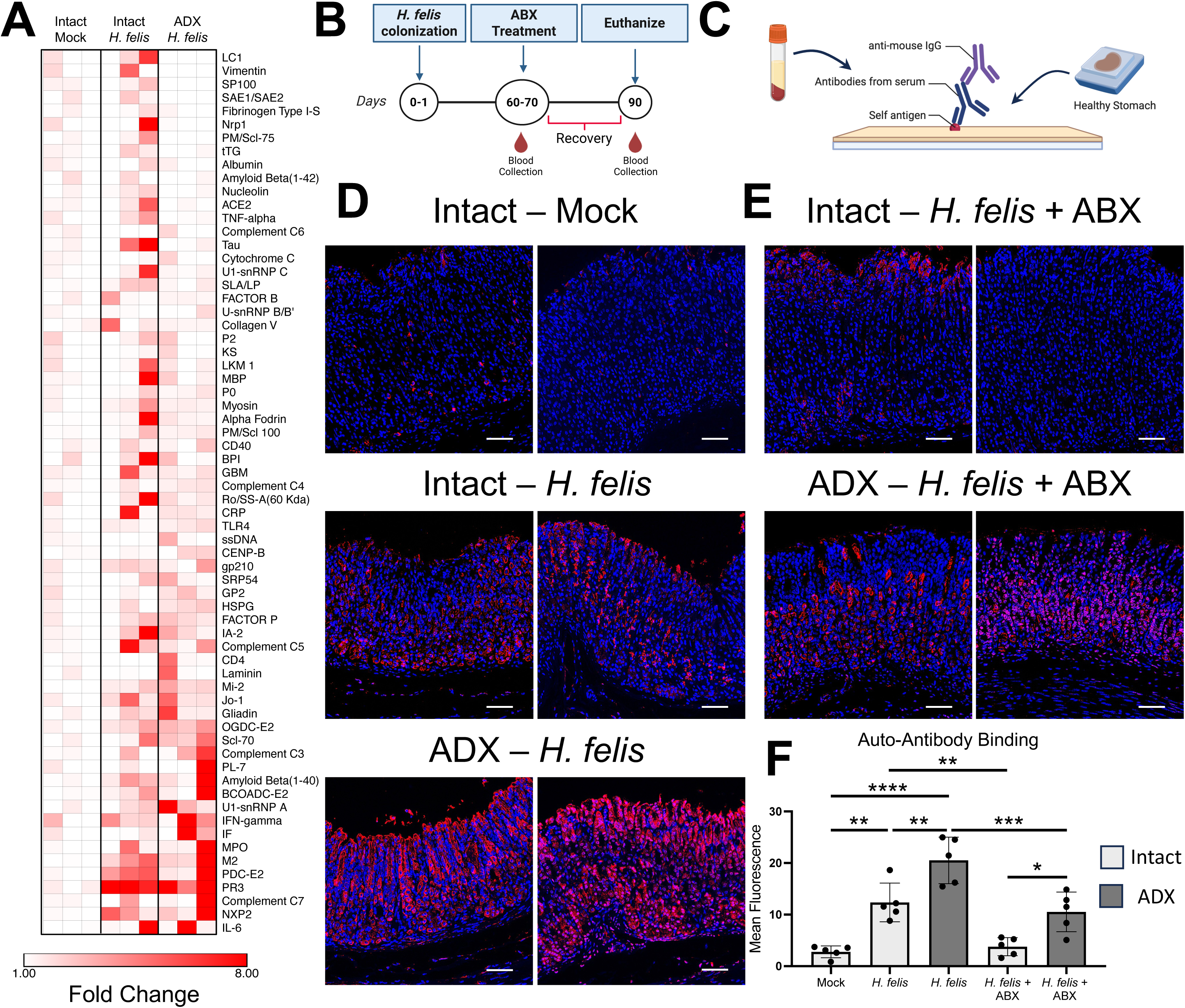
Endogenous glucocorticoids protect from autoimmune reaction in response to *H. felis* colonization. (A) Heatmap of normalized IgG antibody binding scores from serum collected two months post-inoculation. n=3/group. (B-C) Schematics of experimental design visualizing serum collection timeline (B) and gastric autoantibody assay (C). (D-E) Representative immunofluorescent visualizing binding of self-reactive IgG antibodies within the gastric corpus. Pannel D shows binding of serum isolated from intact and ADX mice 2 months after colonization. Pannel E shows binding one month after treatment with antibiotic to eradicate active *H. felis* infection. Two replicate images are shown/group. (F) quantification of mean fluorescence intensity between experimental groups. Scale bars = 65μm. Data are presented as mean ± SD; \**p* < 0.05, **p < 0.01, \*\*\**p* < 0.001, ****p < 0.0001. n=5/group.

## Discussion

Glucocorticoids are well-known immunomodulatory hormones. While they are best known-for their potent immunosuppressive effects at therapeutic levels, endogenous glucocorticoids exert more nuanced and context-dependent influences on immune regulation. We have previously shown that depletion of endogenous glucocorticoids by ADX drives spontaneous gastric inflammation, atrophy, and metaplasia, highlighting their essential role in maintaining immune homeostasis in the stomach ^13–15^. In contrast, endogenous glucocorticoids also promote effective immune responses, and we recently showed that deletion of the GR in the myeloid compartment induced immune dysfunction and ineffective immune responses to *H. pylori* infection ^21^. Thus, endogenous glucocorticoids balance gastric immunity, protecting from pathogenic inflammation by limiting the response to inflammatory stimuli while simultaneously promoting effective responses to *H. pylori* invasion.

*H. pylori* has evolved to evade host immune detection, often persisting for the host’s lifetime unless treated with antibiotics ^22^. In mice, *H. felis* infection closely models this pattern of chronic colonization, and we confirmed that all intact *H. felis-*infected mice remained colonized one-year post-inoculation. In contrast, the majority of ADX mice spontaneously eradicated their *H. felis* infections. Immune evasion is critical for chronic *Helicobacter* colonization. Enhanced Th1 immune responses in mice deficient for the anti-inflammatory cytokine *Il10* leads to rapid *H. felis* eradication ^23^. Similarly, we observed that nearly all ADX mice eradicated their *H. felis* infections, likely resulting from enhanced gastric immune responses and increased expression of Th1-associated cytokines. Th1-dominated immune responses are effective at controlling bacterial infections but also increase host tissue damage. Endogenous glucocorticoids play a critical role in tempering immune activation and directly oppose Th1 responses by inhibiting CD4⁺ T cell activation and downregulating proinflammatory cytokine expression ^12, 24^. Simultaneously, glucocorticoids promote Th2 responses and stimulate the production of anti-inflammatory cytokines such as IL-10. Our findings suggest that *Helicobacter* benefit from the immune suppressive effects of endogenous glucocorticoid signaling, promoting their long-term colonization. Disruption of glucocorticoid signaling undermines this immune evasion strategy, tipping the balance toward immune-mediated clearance.

Immune responses are intricately balanced between tolerance and protection. Tolerant immune responses limit tissue damage and protect from autoimmunity by limiting inflammation and maintaining self-tolerance but increase susceptibility to infections and cancer ^8, 25^. Conversely, hyperactive immune responses may provide more robust protection from infections and tumors but can also damage host tissues and compromise peripheral tolerance, potentially leading to autoimmune disease. In this study, we found that *H. felis* infection promoted autoimmune responses, as all infected mice developed autoantibodies targeting a range of self-antigens. Chronic bacterial infections are well known to elicit autoimmune reactions through mechanisms such as molecular mimicry, epitope spreading, and bystander activation. For instance, antibodies against *S. pyogenes* M protein and N-acetyl-β-D-glucosamine can cross react with heart myosin, leading to rheumatic fever ^26^. Similarly, infection-induced proinflammatory cytokines such as IFNG, TNF, GM-CSF, and IL-6 are associated with activation of self-reactive B and T cells ^6, 27, 28^. Several clinical studies have linked *H. pylori* infection to increased incidence of autoimmune diseases including psoriasis ^29, 30^, type 1 diabetes mellitus ^31–33^, autoimmune thyroiditis ^34, 35^, and systemic lupus erythematosus ^36, 37^. A study of Scandinavian subjects reported anti-*H. pylori* antibodies were detected in 83% of patients with pernicious anemia (also called Addison’s anemia), suggesting a history of prior infection ^38^. Another study reported the presence of anti-parietal cell antibodies in 25% of *H. pylori*-infected individuals without any known autoimmune diseases ^39^. These findings support the notion that, while *H. pylori* infection and AIG are traditionally viewed as distinct diseases, *H. pylori* infection may increase the risk of developing AIG through its capacity to disrupt immune tolerance.

Humoral autoimmunity is common in the context of bacterial infections. However, autoantibody production in this context is typically transient and does not progress to clinical autoimmune disease ^7^. In this study, we found that all *H. felis*-infected mice developed self-reactive IgG antibodies, but these responses subsided following bacterial eradication. This suggests that the autoantibody production was dependent on the inflamed cytokine milieu caused by infection and had not progressed to overt autoimmune disease. In contrast, glucocorticoid depletion enhanced self-reactivity with ADX-infected mice developing IgG antibodies against a broader range of gastric antigens, exhibited higher autoantibody titers, and maintained self-reactive responses even after *H. felis* clearance, suggesting a transition from transient autoreactivity to autoimmune disease. These findings mirror clinical observations in Cushing’s syndrome patients, where a subset of patients develop autoimmune disease following ADX ^40, 41^.

Endogenous glucocorticoids strongly support peripheral tolerance through several mechanisms. They promote the maturation and function of regulatory T cells (Tregs) by inducing the expression of TGFβ, FOXP3, and IL-10 ^42–44^. In our study, ADX mice failed to mount a Treg response during *H. felis* infection, likely exacerbating T cell hyperactivation. Glucocorticoids also support immune tolerance by limiting the production of proinflammatory cytokines and by suppressing antigen presentation and co-stimulatory molecule expression on antigen-presenting cells ^12, 45^. Within sites of inflammation, high levels of proinflammatory cytokines and tissue damage can drive epitope spreading and bystander activation of self-reactive T cells. Both *H. felis* and *H. pylori* induce chronic gastric inflammation, but endogenous glucocorticoids normally oppose this activation by limiting cytokine production and moderating T cell activation. Disruption of glucocorticoid signaling during active *H. pylori* infection would likely exacerbate inflammation and promote autoimmunity. Supporting this notion, we found significantly elevated expression of *Tnf* and *Ifng* in the stomachs of ADX-infected mice, two cytokines that are associated with autoimmune disease progression and are normally suppressed by glucocorticoids ^28, 46, 47^. While glucocorticoids are also involved in thymocyte development—particularly in promoting positive selection—this process is largely independent of adrenal-derived glucocorticoids ^48^. Instead, thymic epithelial cells produce glucocorticoids locally to regulate thymocyte maturation via paracrine signaling. Therefore, the autoimmune responses observed in ADX mice are unlikely to be due to defects in central tolerance and more likely reflect the loss of peripheral regulatory mechanisms governed by adrenal glucocorticoids.

*H. pylori* and autoimmune gastritis are distinct diseases each with separate etiologies ^5^. However, *H. pylori* has long been suspected to promote gastric autoimmunity ^9, 39^. Interestingly, as *H. pylori* rates in the United States continue to decline, there has been an increase in early-onset gastric cancers among women, which has been linked to AIG ^16^. The mechanisms driving this trend remain unclear, although endocrine disruption has been proposed as a contributing factor. In this study, we showed that *H. felis* infection induces autoantibody production, a response that is further amplified by ADX. Moreover, despite clearing their *H. felis* infections, ADX mice exhibited persistent gastric inflammation, with a subset developing dysplasia. These findings suggest that severe inflammation associated with *Helicobacter* infection may simultaneously drive bacterial clearance and promote autoimmune pathology. Thus, individuals with *H. pylori* infection who experience transient disruption of glucocorticoid signaling—whether through exposure to endocrine-disrupting chemicals or treatment with pharmacologic inhibitors such as RU486 ^49^— may face increased risk of developing AIG and a long-term elevated gastric cancer risk.

In summary, the relationship between *H. pylori* infection and autoimmunity remains incompletely understood. While clinical studies have linked *H. pylori* to autoimmune responses, the underlying mechanisms and their clinical implications require further investigation. As *H. pylori* infection rates continue to decline, AIG is predicted to become a leading gastric cancer risk factor in the future ^50^. Our findings provide new evidence that *Helicobacter* infection can trigger autoimmune responses and that disruption of endogenous glucocorticoid signaling increases the intensity of these autoimmune reactions. These data raise the possibility that *H. pylori*-infected individuals who experience impaired glucocorticoid signaling may be particularly susceptible to AIG development. Future studies should further investigate the interplay between *H. pylori* and AIG, as their combined inflammatory effects may significantly alter gastric cancer risk.

## Acknowledgments

This work was supported by West Virginia University start-up funds (J.T.B.) National Institutes of Health grants P20GM121322 (J.T.B.), R01ES031253 (S.H.), R01CA77955 (R.M.P.), R01DK58587 (R.M.P.), P01CA116087 (R.M.P.), P30DK058404 (R.M.P.), and R01CA281732 (R.M.P). M.T.M. received support from the system toxicology training grant (T32ES032920). The West Virginia University Microscope Imaging Facility, Flow Cytometry & Single Cell Core, and Genomics Core Facility receive support from the National Institutes of Health grants P30GM103503 and S10 grant OD028605, and U54 GM104942, respectively.

## Abbreviations

ADX: Adrenalectomy
AIG: adrenal intact (intact) autoimmune gastritis
ABX: Antibiotics
PM: Pyloric Metaplasia

## References

1. Bray F, Ferlay J, Soerjomataram I, Siegel RL, Torre LA, Jemal A. Global cancer statistics 2018: GLOBOCAN estimates of incidence and mortality worldwide for 36 cancers in 185 countries. CA: a cancer journal for clinicians 2018;68:394–424.

2. Rugge M, Bricca L, Guzzinati S, Sacchi D, Pizzi M, Savarino E, Farinati F, Zorzi M, Fassan M, Dei Tos AP, Malfertheiner P, Genta RM, Graham DY. Autoimmune gastritis: long-term natural history in naïve Helicobacter pylori-negative patients. Gut 2023;72:30–38.

3. Chen C, Yang Y, Li P, Hu H. Incidence of Gastric Neoplasms Arising from Autoimmune Metaplastic Atrophic Gastritis: A Systematic Review and Case Reports. J Clin Med 2023;12.

4. Hoft SG, Brennan M, Carrero JA, Jackson NM, Pretorius CA, Bigley TM, Sáenz JB, DiPaolo RJ. Unveiling Cancer-Related Metaplastic Cells in Both Helicobacter pylori Infection and Autoimmune Gastritis. Gastroenterology 2025;168:53–67.

5. Hoft SG, Noto CN, DiPaolo RJ. Two Distinct Etiologies of Gastric Cancer: Infection and Autoimmunity. Front Cell Dev Biol 2021;9:752346.

6. Johnson D, Jiang W. Infectious diseases, autoantibodies, and autoimmunity. J Autoimmun 2023;137:102962.

7. Litwin CM, Binder SR. ANA testing in the presence of acute and chronic infections. J Immunoassay Immunochem 2016;37:439–52.

8. Pacheco Y, Acosta-Ampudia Y, Monsalve DM, Chang C, Gershwin ME, Anaya JM. Bystander activation and autoimmunity. J Autoimmun 2019;103:102301.

9. Neumann WL, Coss E, Rugge M, Genta RM. Autoimmune atrophic gastritis--pathogenesis, pathology and management. Nat Rev Gastroenterol Hepatol 2013;10:529–41.

10. Arnold IC, Lee JY, Amieva MR, Roers A, Flavell RA, Sparwasser T, Müller A. Tolerance rather than immunity protects from Helicobacter pylori-induced gastric preneoplasia. Gastroenterology 2011;140:199–209.

11. Khadka S, Druffner SR, Duncan BC, Busada JT. Glucocorticoid regulation of cancer development and progression. Front Endocrinol (Lausanne) 2023;14:1161768.

12. Cain DW, Cidlowski JA. Immune regulation by glucocorticoids. Nat Rev Immunol 2017;17:233–247.

13. Busada JT, Khadka S, Peterson KN, Druffner SR, Stumpo DJ, Zhou L, Oakley RH, Cidlowski JA, Blackshear PJ. Tristetraprolin Prevents Gastric Metaplasia in Mice by Suppressing Pathogenic Inflammation. Cell Mol Gastroenterol Hepatol 2021;12:1831–1845.

14. Busada JT, Peterson KN, Khadka S, Xu X, Oakley RH, Cook DN, Cidlowski JA. Glucocorticoids and Androgens Protect From Gastric Metaplasia by Suppressing Group 2 Innate Lymphoid Cell Activation. Gastroenterology 2021;161:637–652.e4.

15. Busada JT, Ramamoorthy S, Cain DW, Xu X, Cook DN, Cidlowski JA. Endogenous glucocorticoids prevent gastric metaplasia by suppressing spontaneous inflammation. J Clin Invest 2019;129:1345–1358.

16. Anderson WF, Rabkin CS, Turner N, Fraumeni JF, Jr., Rosenberg PS, Camargo MC. The Changing Face of Noncardia Gastric Cancer Incidence Among US Non-Hispanic Whites. J Natl Cancer Inst 2018;110:608–615.

17. Oh J, Abboud Y, Burch M, Gong J, Waters K, Ghaith J, Jiang Y, Park K, Liu Q, Watson R, Lo SK, Gaddam S. Rising Incidence of Non-Cardia Gastric Cancer among Young Women in the United States, 2000-2018: A Time-Trend Analysis Using the USCS Database. Cancers (Basel) 2023;15.

18. Amieva M, Peek Jr RM. Pathobiology of Helicobacter pylori–induced gastric cancer. Gastroenterology 2016;150:64–78.

19. Bertaux-Skeirik N, Wunderlich M, Teal E, Chakrabarti J, Biesiada J, Mahe M, Sundaram N, Gabre J, Hawkins J, Jian G. CD44 variant isoform 9 emerges in response to injury and contributes to the regeneration of the gastric epithelium. The Journal of pathology 2017;242:463–475.

20. Wada T, Ishimoto T, Seishima R, Tsuchihashi K, Yoshikawa M, Oshima H, Oshima M, Masuko T, Wright NA, Furuhashi S. Functional role of CD 44v-x CT system in the development of spasmolytic polypeptide-expressing metaplasia. Cancer science 2013;104:1323–1329.

21. Khadka S, Dziadowicz SA, Xu X, Wang L, Hu G, Carrero JA, DiPaolo RJ, Busada JT. Endogenous glucocorticoids are required for normal macrophage activation and gastric Helicobacter pylori immunity. Am J Physiol Gastrointest Liver Physiol 2024;327:G531–g544.

22. Sirit IS, Peek RM, Jr. Decoding the Ability of Helicobacter pylori to Evade Immune Recognition and Cause Disease. Cell Mol Gastroenterol Hepatol 2025;19:101470.

23. Ismail HF, Fick P, Zhang J, Lynch RG, Berg DJ. Depletion of neutrophils in IL-10(-/-) mice delays clearance of gastric Helicobacter infection and decreases the Th1 immune response to Helicobacter. J Immunol 2003;170:3782–9.

24. Franchimont D, Galon J, Gadina M, Visconti R, Zhou Y, Aringer M, Frucht DM, Chrousos GP, O’Shea JJ. Inhibition of Th1 immune response by glucocorticoids: dexamethasone selectively inhibits IL-12-induced Stat4 phosphorylation in T lymphocytes. J Immunol 2000;164:1768–74.

25. Yosri M, Dokhan M, Aboagye E, Al Moussawy M, Abdelsamed HA. Mechanisms governing bystander activation of T cells. Front Immunol 2024;15:1465889.

26. Cunningham MW. Molecular Mimicry, Autoimmunity, and Infection: The Cross-Reactive Antigens of Group A Streptococci and their Sequelae. Microbiol Spectr 2019;7.

27. Pollard KM, Cauvi DM, Toomey CB, Morris KV, Kono DH. Interferon-γ and systemic autoimmunity. Discov Med 2013;16:123–31.

28. Suk K, Kim S, Kim YH, Kim KA, Chang I, Yagita H, Shong M, Lee MS. IFN-gamma/TNF-alpha synergism as the final effector in autoimmune diabetes: a key role for STAT1/IFN regulatory factor-1 pathway in pancreatic beta cell death. J Immunol 2001;166:4481–9.

29. Campanati A, Ganzetti G, Martina E, Giannoni M, Gesuita R, Bendia E, Giuliodori K, Sandroni L, Offidani A. Helicobacter pylori infection in psoriasis: results of a clinical study and review of the literature. Int J Dermatol 2015;54:e109–14.

30. Onsun N, Arda Ulusal H, Su O, Beycan I, Biyik Ozkaya D, Senocak M. Impact of Helicobacter pylori infection on severity of psoriasis and response to treatment. Eur J Dermatol 2012;22:117–20.

31. Zekry OA, Abd Elwahid HA. The association between Helicobacter pylori infection, type 1 diabetes mellitus, and autoimmune thyroiditis. J Egypt Public Health Assoc 2013;88:143–7.

32. Bazmamoun H, Rafeey M, Nikpouri M, Ghergherehchi R. Helicobacter Pylori Infection in Children with Type 1 Diabetes Mellitus: A Case-Control Study. J Res Health Sci 2016;16:68–71.

33. Rahman MA, Cope MB, Sarker SA, Garvey WT, Chaudhury HS, Khaled MA. Helicobacter pylori Infection and Inflammation: Implication for the Pathophysiology of Diabetes and Coronary Heart Disease in Asian Indians. J Life Sci 2009;1:45–50.

34. Hou Y, Sun W, Zhang C, Wang T, Guo X, Wu L, Qin L, Liu T. Meta-analysis of the correlation between Helicobacter pylori infection and autoimmune thyroid diseases. Oncotarget 2017;8:115691–115700.

35. Bertalot G, Montresor G, Tampieri M, Spasiano A, Pedroni M, Milanesi B, Favret M, Manca N, Negrini R. Decrease in thyroid autoantibodies after eradication of Helicobacter pylori infection. Clin Endocrinol (Oxf) 2004;61:650–2.

36. Youssefi M, Tafaghodi M, Farsiani H, Ghazvini K, Keikha M. Helicobacter pylori infection and autoimmune diseases; Is there an association with systemic lupus erythematosus, rheumatoid arthritis, autoimmune atrophy gastritis and autoimmune pancreatitis? A systematic review and meta-analysis study. J Microbiol Immunol Infect 2021;54:359–369.

37. Wu MC, Leong PY, Chiou JY, Chen HH, Huang JY, Wei JC. Increased Risk of Systemic Lupus Erythematosus in Patients With Helicobacter pylori Infection: A Nationwide Population-Based Cohort Study. Front Med (Lausanne) 2019;6:330.

38. Ma JY, Borch K, Sjöstrand SE, Janzon L, Mårdh S. Positive correlation between H,K-adenosine triphosphatase autoantibodies and Helicobacter pylori antibodies in patients with pernicious anemia. Scand J Gastroenterol 1994;29:961–5.

39. Claeys D, Faller G, Appelmelk BJ, Negrini R, Kirchner T. The gastric H+,K+-ATPase is a major autoantigen in chronic Helicobacter pylori gastritis with body mucosa atrophy. Gastroenterology 1998;115:340–7.

40. Petramala L, Olmati F, Conforti MG, Concistré A, Bisogni V, Alfieri N, Iannucci G, de Toma G, Letizia C. Autoimmune Diseases in Patients with Cushing’s Syndrome after Resolution of Hypercortisolism: Case Reports and Literature Review. Int J Endocrinol 2018;2018:1464967.

41. Takasu N, Komiya I, Nagasawa Y, Asawa T, Yamada T. Exacerbation of autoimmune thyroid dysfunction after unilateral adrenalectomy in patients with Cushing’s syndrome due to an adrenocortical adenoma. N Engl J Med 1990;322:1708–12.

42. Karagiannidis C, Akdis M, Holopainen P, Woolley NJ, Hense G, Rückert B, Mantel PY, Menz G, Akdis CA, Blaser K, Schmidt-Weber CB. Glucocorticoids upregulate FOXP3 expression and regulatory T cells in asthma. J Allergy Clin Immunol 2004;114:1425–33.

43. Bereshchenko O, Coppo M, Bruscoli S, Biagioli M, Cimino M, Frammartino T, Sorcini D, Venanzi A, Di Sante M, Riccardi C. GILZ promotes production of peripherally induced Treg cells and mediates the crosstalk between glucocorticoids and TGF-β signaling. Cell Rep 2014;7:464–475.

44. Gayo A, Mozo L, Suárez A, Tuñon A, Lahoz C, Gutiérrez C. Glucocorticoids increase IL-10 expression in multiple sclerosis patients with acute relapse. J Neuroimmunol 1998;85:122–30.

45. Chamorro S, García-Vallejo JJ, Unger WW, Fernandes RJ, Bruijns SC, Laban S, Roep BO, t Hart BA, van Kooyk Y. TLR triggering on tolerogenic dendritic cells results in TLR2 up-regulation and a reduced proinflammatory immune program. J Immunol 2009;183:2984–94.

46. Hu X, Li WP, Meng C, Ivashkiv LB. Inhibition of IFN-gamma signaling by glucocorticoids. J Immunol 2003;170:4833–9.

47. Steer JH, Kroeger KM, Abraham LJ, Joyce DA. Glucocorticoids suppress tumor necrosis factor-alpha expression by human monocytic THP-1 cells by suppressing transactivation through adjacent NF-kappa B and c-Jun-activating transcription factor-2 binding sites in the promoter. J Biol Chem 2000;275:18432–40.

48. Taves MD, Ashwell JD. Glucocorticoids in T cell development, differentiation and function. Nat Rev Immunol 2021;21:233–243.

49. Morris MT, Pascoe JL, Busada JT. In vitro to in vivo evidence for chemical disruption of glucocorticoid receptor signaling. Toxicol Rep 2025;14:102053.

50. Song M, Rabkin CS, Camargo MC. Gastric Cancer: an Evolving Disease. Curr Treat Options Gastroenterol 2018;16:561–569.

